# Forecasting the global extent of invasion of the cereal pest *Spodoptera frugiperda*, the fall armyworm

**DOI:** 10.1101/391847

**Authors:** Regan Early, Pablo González-Moreno, Sean T. Murphy, Roger Day

## Abstract

Fall armyworm, *Spodoptera frugiperda*, is a crop pest native to the Americas, which has invaded and spread throughout sub-Saharan Africa within two years. Recent estimates of 20-50% maize yield loss in Africa suggest severe damage to livelihoods. Fall armyworm is still infilling its potential range in Africa, and could spread to other continents. In order to understand fall armyworm’s year-round, global, potential distribution, we used evidence of the effects of temperature and precipitation on fall armyworm life-history, combined with data on native and African distributions to construct Species Distribution Models (SDMs). Fall armyworm has only invaded areas that have a climate similar to the native distribution, validating the use of climatic SDMs. The strongest climatic limits on fall armyworm’s year-round distribution are the coldest annual temperature and the amount of rain in the wet season. Much of sub-Saharan Africa can host year-round fall armyworm populations, but the likelihoods of colonising North Africa and seasonal migrations into Europe are hard to predict. South and Southeast Asia and Australia have climate that would permit fall armyworm to invade. Current trade and transportation routes reveal Australia, China, India, Indonesia, Malaysia, Philippines, and Thailand face high threat of fall armyworm invasions originating from Africa.

## Introduction

Fall armyworm, *Spodoptera frugiperda* (J.E. Smith) (Lepidoptera: Noctuidae) is native to the Americas, from as far south as La Pampa, Argentina, to as far north as southern Florida and Texas, USA. Fall armyworm caterpillars are major pests of cereals and forage grasses, and recorded as eating 186 plant species from 42 families^1^. In Florida, fall armyworm is the most serious pest of maize, causing up to 20% yield loss. In areas where less money is available for pest management, impacts are even more severe. For example, yield losses can reach 40% in Honduras^2^, and 72% in Argentina^3^. Other plants that are economically important in Africa and that suffer major fall armyworm damage include rice, sugarcane, sorghum, beet, tomato, potato, cotton, and pasture grasses^4,5^.

In January 2016 major outbreaks of armyworms were reported in South West Nigeria, and Ghana, and shortly after in Benin, Sao Tomé, and Togo^6^. Morphological and molecular analysis by IITA found that the armyworms were *S. frugiperda*, and not the native armyworms *S. exigua* or *exempta*^7^. As of 28^th^ September 2017 28 sub-Saharan African countries had confirmed presence of fall armyworm, with nine more suspecting or awaiting confirmation of the species’ presence^4^. Within these countries the fall armyworm is still spreading^8^. The countries of the initial outbreaks (Ghana, Benin, Togo, and Nigeria) have West Africa’s major air transportation hubs, and are highly climatically similar to the regions from which arriving flights originate^9,10^. As a result this region is one of the areas most likely to act as an epicentre for invasions in Africa^11^. Indeed, it is speculated that fall armyworm entered Africa as a stowaway on a passenger flight^12^; unaided dispersal is considered unlikely because prevailing winds are generally from East to West. Molecular data from specimens in Togo indicate that fall armyworm in Africa originates from an area encompassing the eastern USA, Caribbean, and Lesser Antilles^13^. The two known strains of the species, the so-called maize- and rice strains, overlap in the latter region and both strains have been found in Africa (Cock *et al*. 2017, Nagoshi *et al*. 2017). Considering the host and time mating differences between the strains^14^ the likelihood of both strains to be introduced together is rather low. Thus, it is still unclear if the invasion has originated from a single introduction or multiple from the same region.

The rapid spread throughout Africa was likely due to adult fall armyworm’s ability to travel very long distances. Adults can travel several hundred kilometres in a single night by flying to and maintaining an elevation of several hundred metres, at which height winds can transport adults in a directional manner^15^. Thus, from its current known distribution fall armyworm could appear at almost any point in its potential African range literally overnight. Furthermore, fall armyworm’s long distance dispersal ability means that it can undergo seasonal migrations of thousands of kilometres. Thus, although Fall armyworm cannot survive cold temperatures by entering diapause, it still poses a threat to temperate regions^16^. Between spring and autumn three successive generations of fall armyworm travel 1700km north from Texas and Florida to infest crops as far north as Québec and Ontario^15^. Adults also appear to migrate several hundred kilometres over the sea^15^. This means that North African countries could be within the reach of fall armyworm from sub-Saharan populations. If the species can survive year-round in North Africa, then seasonal migrations into Europe could also occur. This would pose a severe threat to agriculture in Europe, and the fall armyworm is classed as a Quarantine Pest for the EU^17^. Further afield, the fall armyworm’s wide distribution in the Americas and Africa suggest that it could establish easily in East and Southeast Asia. Given increasing levels of trade and transportation between infested parts of Africa and the rest of the world^10^, it seems likely that fall armyworm could be unwittingly transported onwards to environmentally suitable regions.

Given fall armyworm’s potentially large global range, countries and regions that depend on crops the fall armyworm threatens urgently need information on the pest’s likely potential distribution and environmental limitations. Such information would inform national and regional pest risk assessments and appropriate management strategies in several ways: by quantifying the agricultural areas within Africa that are at risk from year-round populations or seasonal migrations, informing the likelihood of seasonal migrations into Europe, and classifying the likelihood of establishment if fall armyworm is transported into other parts of the world. This information would also help target awareness raising and monitoring for early detection. Early detection is essential as fall armyworm can currently only be effectively controlled with chemical insecticides while the larvae are small^18^. Early-stage larvae live inside the whorl of the developing plant, and so are hard to detect with casual observation. Detecting fall armyworm in time to apply pesticide and avoid heavy crop losses means inspecting plants for eggs or larvae, or setting pheromone traps for adults^8^. Raising awareness amongst farmers and developing surveillance schemes is therefore a priority for managers, particularly in Africa^8^. Knowing where outbreaks are likely to occur would encourage and inform these efforts.

Here we first reviewed what is known of the environmental controls on the fall armyworm’s lifecycle and herbivory, particularly on maize, the crop most economically important and threatened by the moth in Africa^8^. We then quantified the species’ environmental limitations and potential distribution worldwide, using information on the native and invasive distribution to construct an ensemble SDM. We assessed the robustness of this approach by asking whether fall armyworm has invaded parts of Africa a climatic SDM would not predict, i.e. undergone a ‘niche shift’^19^. The SDMs we constructed are based on annual conditions, and thus predict suitability for year-round populations of fall armyworm. Lastly, we use trade and transportation links to interpret the potential for the fall armyworm to spread beyond Africa via these pathways.

The accuracy of SDM results and the similarity of the environments occupied in the native and invaded range support the robustness of the SDM approach. The temperature of the coldest month and the amount of rain during the rainy season are the most important climatic limits of fall armyworm’s year-round distribution. Much of sub-Saharan Africa can host year-round fall armyworm populations, and seasonal migrations are likely to take place along the Nile into Northeast Africa. The likelihood of seasonal migrations beyond this range seems to be low. South and Southeast Asia, and Australia, are highly suitable for fall armyworm. Trade and passenger air travel routes indicate parts of this region into which African populations are particularly likely to be transported.

## Results

### Has the fall armyworm invaded where we expect it to, based on the native distribution in the Americas?

All but 3% of the recorded current distribution in Ghana and Zambia is found in climate and forest conditions that match the native range, i.e. there is virtually no orange area in Fig. 3. Fall armyworm therefore does not appear to have undergone niche shift during invasion.

**Figure 1.**
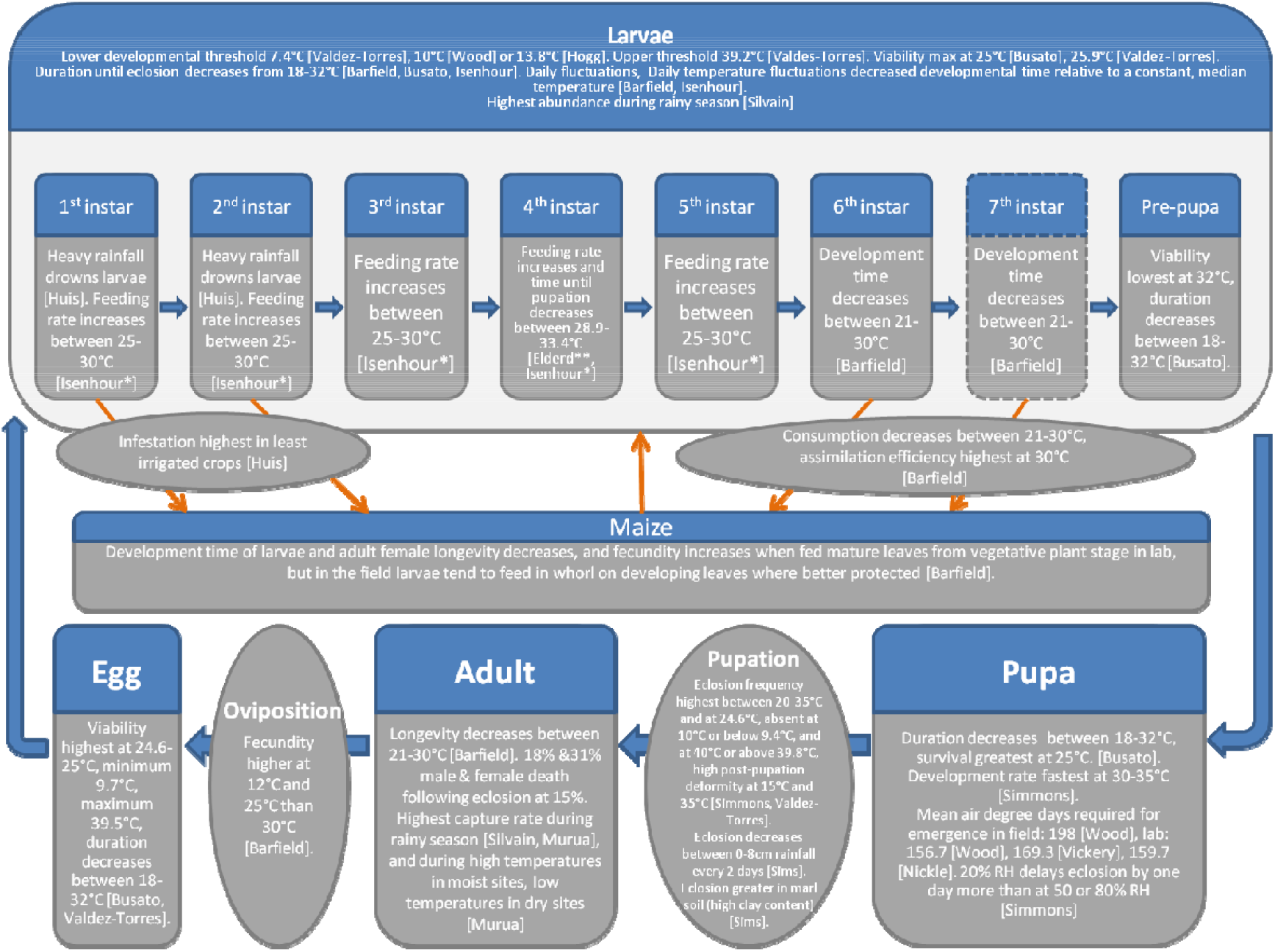
Empirically measured environmental effects on fall armyworm life cycle. Summary of data from literature of temperature, moisture, soil, and host plant effects on fall armyworm survival and developmental time, and observations of abundances in the field under different conditions and seasons. All studies were conducted with populations from the Americas. Rectangles are life stages, ovals are processes. Orange arrows represent effects of fall armyworm on maize or vice versa. The occurrence of a 7^th^ instar is not universally reported. Only direct effects (measured or imputed) of the environment and host on fall armyworm were included. Unless otherwise noted, moths for experiments were reared on fresh maize or an artificial diet. *Feeding rate applies to first nine days of larval stages, which is approximately the first four larval instars. **Feeding rate measured on soybean leaf.

**Figure 2.**
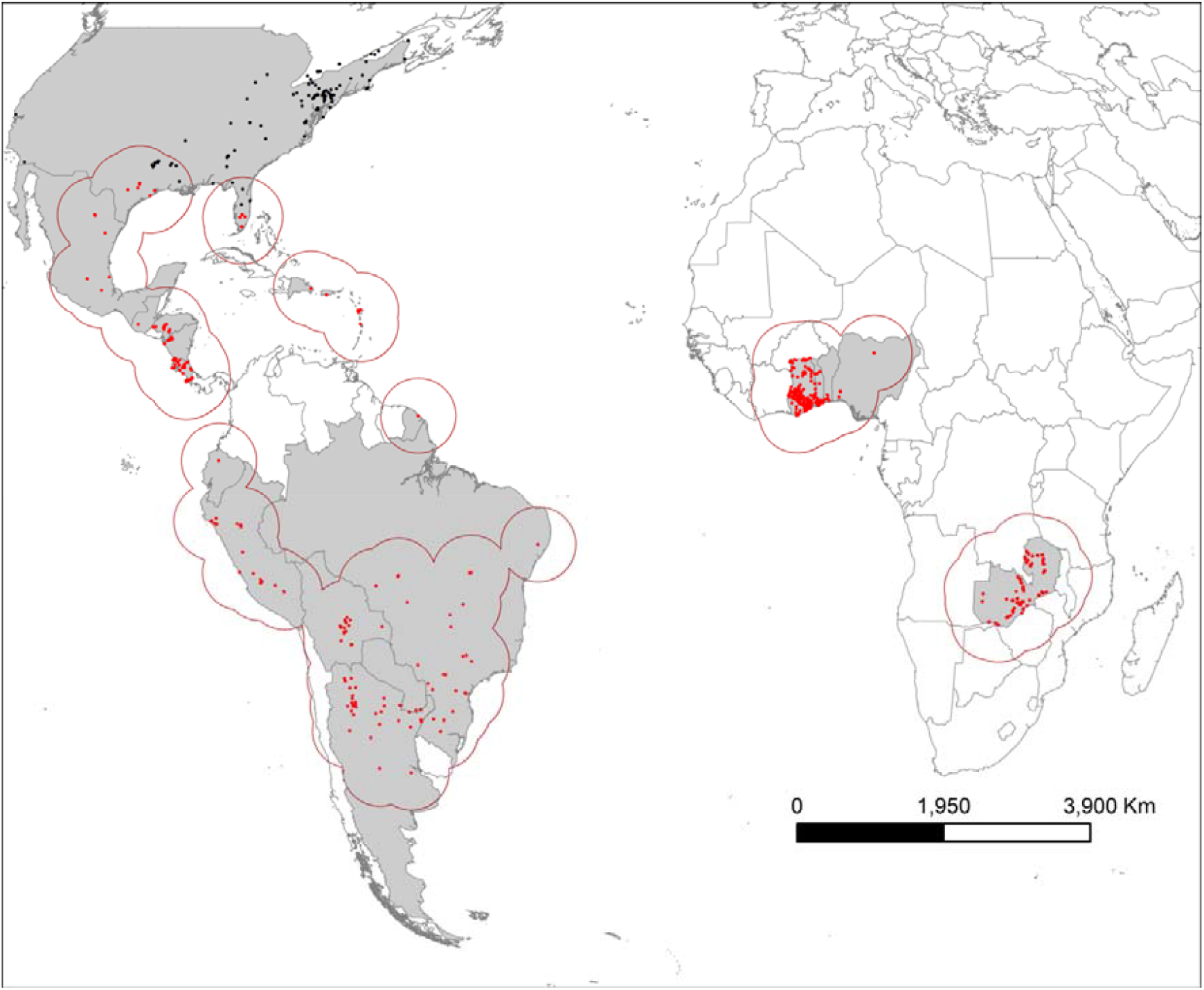
Fall armyworm distribution data. Red points are the complete set of presence locations used to make models. Black points in the USA are not part of the year-round native distribution and were not included in models. Grey areas are the geographically unrestricted background from which pseudo-absences could be drawn. Red lines indicate the geographically restricted background (500km radius around presence locations) within which pseudo-absences could be drawn.

**Figure 3.**
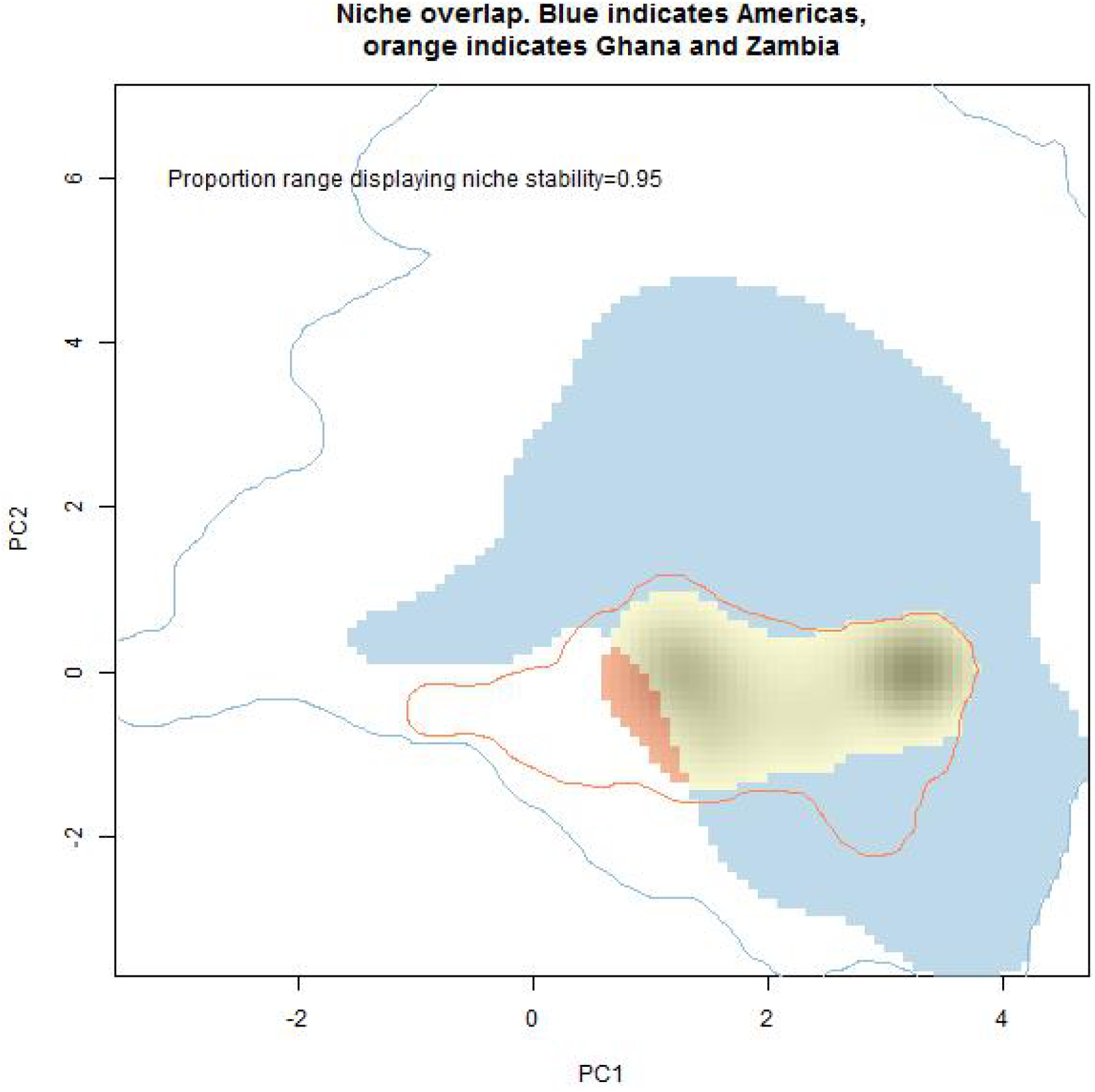
Similarity of environmental niches in the Americas and Africa. The outlines represent the environmental conditions available in the Americas (blue), and Ghana and Zambia (orange). The blue shaded area represents the conditions the species occupies only in the Americas. The yellow and orange shaded areas represent the conditions the species occupies in Ghana and Zambia. The orange area shows the part of the African range that is found in different environmental conditions to the native range (expansion), while the yellow area indicates common conditions (stability). The shading under the yellow/orange area is the density of the species’ occurrence in Ghana and Zambia.

### SDM performance

Internal cross-validation indicated that TSS scores were ‘moderate’ and AUC scores were ‘fair’ (Table 1). These scores increased as more of the African distribution data were included in analyses. The Spearman correlations between global projections made using the validation data (70% of each distribution data set) and all data in each distribution dataset were ≥0.87. This indicates that the TSS and AUC scores from internal validation reflect the accuracy of SDMs calculated with all data in each dataset.

**Table 1.** Summary statistics for Species Distribution Models (SDMs). Dataset indicates the percentage of the African distribution data that were sub-sampled. TSS is True Skill Statistic. AUC is Area Under the Receiver Operating Curve. Both indicate predictive accuracy. Mean (±standard deviation) TSS and AUC values are averages calculated by internal cross-validation for all SDMs constructed with each dataset, excluding SDMs that were discarded in making the ensemble due to low predictive accuracy (TSS<0.4). TSS values between 0.4 and 0.5 are often considered ‘moderate’^52^. AUC values between 0.7 and 0.8 are often considered “fair”. Spearman’s ρ is calculated between the global ensemble projections made using 70% (used for internal crossvalidation) and 100% of the data, and indicates whether the validation statistics can be considered representative of the final model. Kappa measures the agreement between the global ensemble forecast using 100% of the distribution data and the entire background and the named dataset. Kappa values of 0.4-0.6 are considered ‘moderate’, and 0.6-0.8 ‘substantial’^52^.

| Pseudo-absence restriction radius | Dataset | TSS | AUC | Spearman’s $\rho$ | Kappa |
| --- | --- | --- | --- | --- | --- |
| None | 5% | 0.52±0.06 | 0.79±0.05 | 0.88±0.12 | 0.49 |
| None | 10% | 0.52±0.07 | 0.80±0.05 | 0.87±0.15 | 0.50 |
| None | 20% | 0.54±0.09 | 0.80±0.06 | 0.89±0.09 | 0.57 |
| None | 30% | 0.55±0.06 | 0.80±0.04 | 0.87±0.12 | 0.59 |
| None | 50% | 0.51±0.07 | 0.79±0.05 | 0.89±0.11 | 0.75 |
| None | 70% | 0.55±0.07 | 0.91±0.05 | 0.91±0.09 | 0.76 |
| None | 100% | 0.55±0.07 | 0.81±0.06 | 0.89±0.08 | NA |
| 500km | 100% | 0.48±0.07 | 0.78±0.04 | 0.79±0.23 | 0.41 |

Agreement between the ensemble projections resulting from sub-sampled and complete datasets was ‘moderate’ to ‘substantial’ (Cohen’s kappa values Table 1, Supplementary Fig. S1). Agreement increased with the percentage of the African data sub-sampled (Supplementary Fig. S2). Ensemble projections from different datasets agreed most strongly in the areas that are least suitable, i.e. <0.2 (Supplementary Fig. S2). The proportion of grid-cells classified within the same suitability band (‘balanced accuracy’) by different ensembles was never less than 0.6 when pseudo-absences were selected from the entire geographic background, and increased between ensembles constructed with similar degrees of sub-sampling (Supplementary Fig. S2).

The dataset using 100% of the data and the entire geographic background gave SDMs that had the highest AUC and TSS scores (Table 1). There was very little difference in the shape or extent of the area in the native American region predicted to be suitable using different sub-samples of the distribution data (Supplementary Fig. S1). This, and the high degree of agreement in the global projections from SDMs using all and sub-sampled African data (Figs S1, S2) suggest that SDMs using 100% of the data did not appear to be over-fit to the African distribution data. SDMs constructed using pseudo-absences drawn from a 500m buffer around presence points appeared to under-predict both the American and African distribution (Fig. 4), and had the lowest TSS and AUC scores (Table 1). We therefore saw no reason not to use predictions from SDMs using the complete distribution dataset and the entire geographic background to represent fall armyworm’s global potential distribution. These SDMs were used in the final ensemble global projection (Fig. 4).

**Figure 4.**
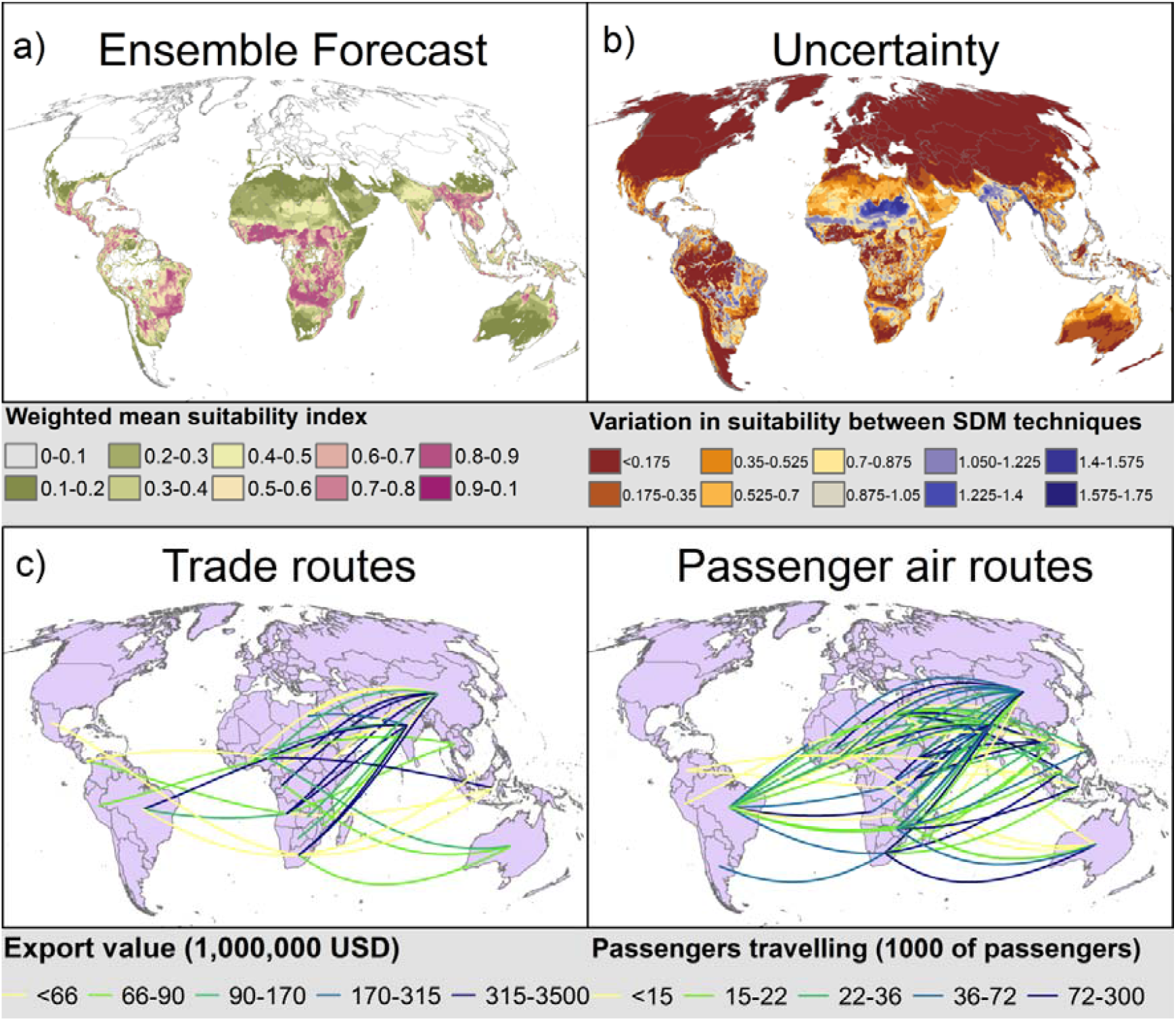
a) Potential global distribution of fall armyworm, as predicted by an ensemble of SDMs constructed using all distribution data and with four pseudo-absence datasets. SDMs were permitted into the ensemble if the TSS from internal cross-validation was ≥0.4. The ensemble was calculated as the mean of projections from all permitted SDMs, each model weighted by the crossvalidated TSS. b) Uncertainty in projections, as calculated by the variation between all projections included in the ensemble. c) value of all exports from 2012-2016 from source sub-Saharan African countries climate to vulnerable countries outside sub-Saharan Africa. The top 5% of trading relationships between these countries are shown, and the five colour categories represent 20% quantiles of export values. d) number of passengers in 2013 travelling from source sub-Saharan African countries with their final destination in vulnerable countries outside sub-Saharan Africa. The top 13% of travel routes between these countries are shown, and the five colour categories represent 20% quantiles of export values.

MinTemp was the most important environmental variable, followed by forest and SumWet (Fig. 5). SeasPpn and LenWet were relatively unimportant. The degree of sub-sampling did not affect variable importance. The range of the environmental conditions from which fall armyworm is recorded is shown in Supplementary Fig. S3.

**Figure 5.**
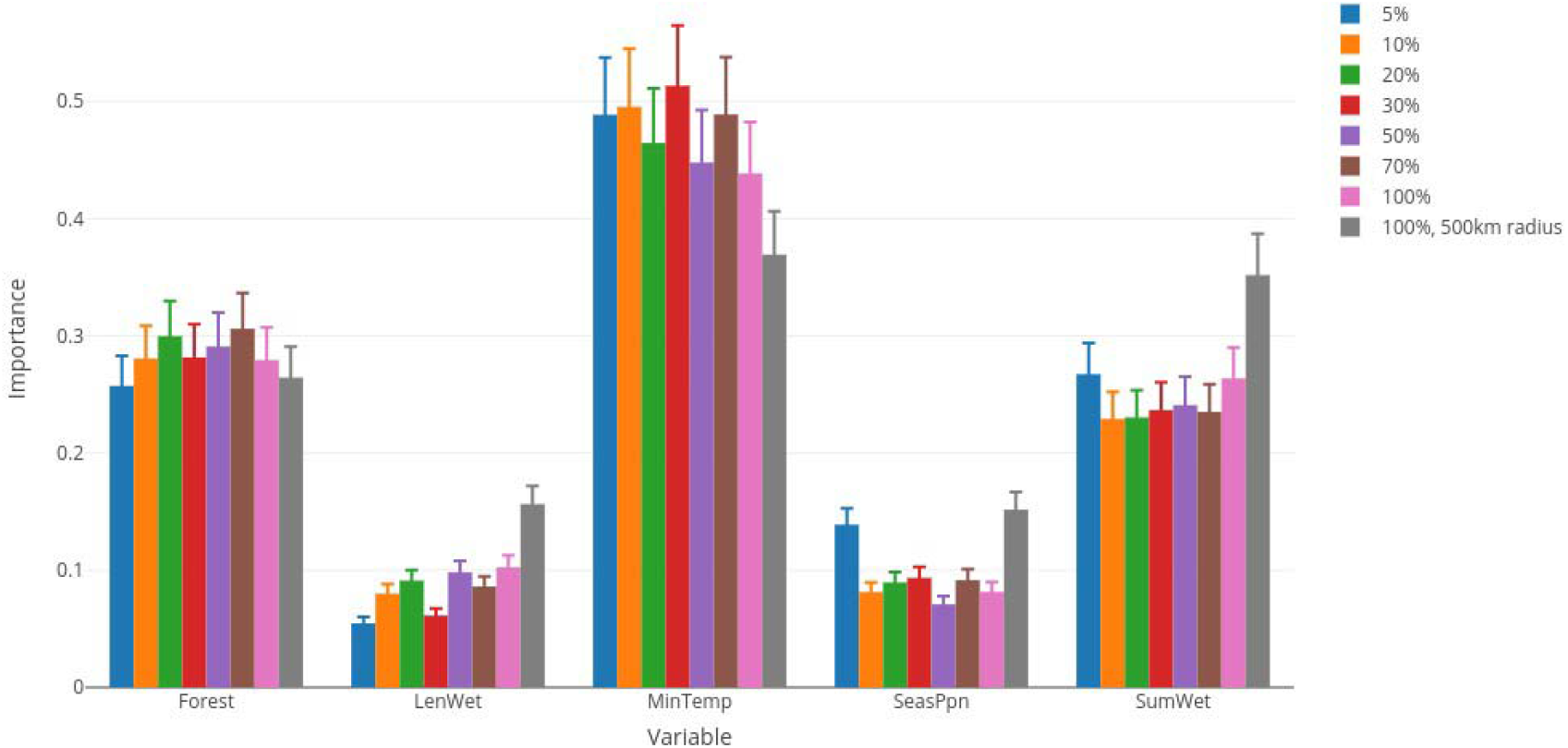
Importance of variables for Species Distribution Models (SDMs) of fall armyworm in the Americas and Africa. Colour codes indicate the percentage of the African distribution that was subsampled or the pseudo-absence selection background. Error bars are standard deviations of the results across all SDM techniques and distribution datasets.

MESS indicated very few areas in which environmental conditions had no analogue in the training region (Supplementary Fig. S4).

### Transportation beyond Africa

Countries vulnerable to fall armyworm (outside South America) that receive the greatest value of commodities exported from African fall armyworm source countries are China, India Indonesia, and to a lesser extent Australia and Thailand. Countries vulnerable to fall armyworm (outside South America) that receive the greatest number of passengers embarking from African source countries are Australia, China, India, Indonesia, Malaysia, and the Philippines. These countries are likely to be the most imminently threatened by fall armyworm invasion.

## Discussion

Species Distribution Modelling indicates that much of sub-Saharan Africa is highly suitable year-round for fall armyworm, from the Saharan belt to South Africa. Within this region, much of Congo, DRC, Gabon, and Cameroon have low suitability (though uncertainty is high in some of these areas). Low suitability in these countries is likely because of extensive forest cover. However, this does not mean that pockets of suitable habitat in those countries will not be severely affected, given the ability of fall armyworm to travel long distances (see below for further discussion of forested areas).

In addition to Ghana and Zambia, fall armyworm’s distribution has been well characterised in South Africa. These data were not available to include in models, but visual inspection demonstrates that SDMs seem to predict the extent of fall armyworm’s distribution in South Africa as reported in January 2017 (<u>http://www.grainsa.co.za/the-invasion-of-the-fall-armyworm-in-south-africa</u>).

Much of Northwest and Northeast Africa has low suitability (<40% probability of occurrence, Fig. 4), so might not host year-round populations of fall armyworm. However, Sudan’s and Egypt’s Nile Valley may be suitable during wet parts of the year, and is adjacent to fall armyworm’s likely year-round range in South Sudan and Ethiopia. Fall armyworm’s 1700km annual migration in North America suggests a similar migration is possible into the Nile Valley, which could threaten maize and cotton production. Indeed, migrating Fall armyworms are a severe pest of cotton in the USA^20^. Simulations of fall armyworm dispersal from the Khartoum area of Sudan and the Addis Ababa area of Ethiopia confirm this as a strong possibility^21^.

In currently un-invaded portions of Africa, there are pockets of high suitability in Morocco’s productive agricultural regions, as well as the Libyan coast. Transportation to North Africa via trade or air transportation routes from sub-Saharan Africa is relatively unlikely (Fig. 4). The suitable areas in Morocco are 2,500km straight-line distance from the year-round distribution. This is far beyond the distance travelled by individuals from a single fall armyworm population in North America, and so colonisation directly from the year-round distribution may be unlikely. However, some Lepidoptera are thought to migrate across the Sahara annually^22^. If fall armyworm were to establish in Morocco, seasonal migrations into Europe would be highly likely. There are pockets of climate suitable for year-round populations (i.e. grid-cells with a suitability value of ≥0.5) in south and northeast Iberia, Italy, and Greece.

Nagoshi, et al.^23^ suggest that fall armyworm movement within Central and South America could occur in response to seasonal changes in rainfall, temperature, and agricultural plantings. Dispersal within this region (i.e. not just the long-distance North American seasonal migration) could be considerable, given genetic mixing amongst populations as widely dispersed as Argentina, Mexico, Mississippi^24^. Therefore, low-suitability areas in sub-Saharan Africa may still experience infestation from migrating fall armyworm during some seasons. This would make it very difficult to control fall armyworm outbreaks by managing any single location. Instead management would have to be coordinated across regions and country borders. Future research into seasonal migration and population dynamics within sub-Saharan Africa is needed.

Understanding the potential for annual migrations both within and beyond of the year-round range requires forecasts of the speed, direction, and heights of prevailing wind during periods when fall armyworm populations are large. We also need to know the migration capacity of African fall armyworm populations. The propensity of individuals to migrate, and the length of time for which adults can fly varies within populations, often genetically^25^. Migration capacity in Africa may differ from that in the putative native range, due to founder effects and selection during introduction.

Much of South and Southeast Asia and areas of Australia are highly suitable for fall armyworm year-round, and there are important invasion routes from Africa into several countries in the region (Fig. 4). An additional possible pathway for invasion into South Asia is the southwest monsoon that blows from Africa to India every year from about June, though further research is needed. The potential for onward fall armyworm invasion is graphically illustrated by the invasion of the South American tomato leafminer, *Tuta absoluta* (Meyrick) (Lepidoptera: Gelechiidae). The pest was first observed outside South America in Spain in 2006, most likely having originated from one source close to or in Chile^26^. It colonised the Middle East and Northeast Africa around 2012, and parts of sub-Saharan Africa from about 2014^27^. Later in 2014 the leafminer was recorded from India^28^ and subsequently invaded other countries in Asia.

Arrival *via* imported commodities or passenger air travel is most likely where entry ports are found in environmentally suitable locations for a given pest^29^. Targeted screening and rapid response mechanisms could help reduce the likelihood of arrival and establishment in these locations. However, management efforts should not be confined to individual countries. There are large, spatially cohesive, areas of environmentally suitable areas throughout Asia and Australia (Fig. 4a). The rapid spread in Africa suggests that if fall armyworm reaches one location in Asia or Australia it could spread throughout the entire region using its own dispersal mechanisms, rather than simply establishing near to the arrival point. Thus, if fall armyworm arrives anywhere in Asia or Australia, rapid cross-border communication and collaboration will be key to effective management.

As conditions outside the predicted year-round range (for example Europe) might be suitable in certain seasons, improved predictions of seasonal suitability could be achieved with demographic modelling of data from lab or field trials, or from statistical modelling e.g. ^30^. If demonstrated to be robust, demographic predictions could identify areas where crops eaten by fall armyworm are grown during seasons when migration could occur.

The results of Species Distribution Modelling were encouragingly accurate. AUC values from crossvalidation were well within the range usually considered acceptable for SDM studies of invertebrates^31^. This is encouraging given the somewhat uneven recording effort, the likelihood that fall armyworm is under-recorded in the Americas, and the large area for which projections were made. SDM projections were consistent across sub-sampled data. Indeed, areas in Colombia, Panama, Venezuela, and Brazil known to harbour fall armyworm but from which no or few presences were recorded were predicted to be environmentally suitable (Fig. 4). Projections from ensembles using differing data sub-samples did not lead to substantially lower suitability or greatly differing geographic patterns of suitability (Figs. S1, S2). This indicates that the intensive surveying in Ghana and Zambia has not caused over-fitting to environmental conditions in those countries.

Forest, MinTemp (coldest annual temperature), and SumWet (rainfall during the wettest three months) were consistently identified as the environmental variables that most affected fall armyworm’s distribution. Fall armyworm is most commonly found in areas with very little forest cover, a minimum annual temperature of 18-26°C, and with 500-700mm rainfall in the three wettest months (Supplementary Fig. S3). The importance of MinTemp supports the existence of a hard poleward geographical boundary, caused by one or more months where temperature drops below a threshold. SumWet was consistently important than LenWet (rainy season length) or SeasPpn (the contrast between the rainy and dry seasons). In order to understand if this is due to indirect (i.e. through host plant growth) or direct effects, one could use structural equation modelling incorporating the yield of key host plants, or incorporate life-history parameters from Supplementary Table S1 into demographic models. The importance of ‘Forest’ is likely because it indicates the availability of crops on which fall armyworm feeds. This could also suggest that few people have looked for fall armyworm outside of areas with extensive crop coverage. It would be useful to study fall armyworm’s survival in forested habitat, and in small cropped areas surrounded by forest. This would inform models of invasion into Congo, DRC, Gabon, and Cameroon, where low projected suitability is likely caused by high forest cover.

Rapid evolution of climate tolerances can occur in pest insects following invasion, i.e. a ‘niche shift’^32,33^. Evolutionary niche shift would compromise invasion forecasts using any modelling technique relying on data from native populations. This includes SDMs^19^, CLIMEX^34^, and physiological and demographic approaches such as Insect Life Cycle Modeller^30^. In addition to evolution, niche shift can also occur during invasion due to species’ native distributions not occupying all environmentally suitable locations. However, non-evolutionary niche shift is uncommon in widespread agricultural pests such as fall armyworm^19^. In any case, niche shift analysis finds that rapid evolution of climate tolerances does not seem to have occurred for the fall armyworm (Fig. 3). It therefore appears that we can be confident in the accuracy of range forecasts that utilise native distribution data. However, there is still a significant part of the niche unoccupied in Ghana and Zambia that is suitable for the species based on the native range distribution. Thus, it is likely that the species will continue its rapid infilling of its potential African range^5^.

It is interesting to note that fall armyworm was not high on a recent list of pest species likely to invade West Africa (in the lower 50th percentile for Ghana, Nigeria, and Togo^35^, invasion rankings obtained from personal communication with Dean Paini). This list was constructed using a Self-Organising Map (SOM) approach. SOM calculates the relative likelihood a target pest species will establish in a region based on whether pests whose range overlaps the target species have also invaded there. The low establishment index calculated for fall armyworm is likely because few pest species have previously jumped from the Americas to Africa, which in turn is presumably because of historically low trade between the two continents^12,36^. While the SOM approach is highly valuable, rapidly changing global trade and transportation patterns open invasion routes that pest species rarely travelled in the past^11,36^. Therefore there may be many other inter-continental pest invasions that are hard to predict.

Very little research has been done into differing climate tolerances between maize and rice strains, and there is insufficient information on their respective distributions to apply SDMs to each strain. Slight differences in basal temperatures of the two strains result in approximately one more generation of the rice strain per year at the optimum temperature of 25°C, i.e. 12 generations^37^. The two strains appear to have substantially overlapping ranges, so any difference in tolerances is likely not to affect range greatly, but could affect abundance and impact. The African maize strain population appears to be a single haplotype^4^. This haplotype may have slightly more restricted climate tolerances than the species as a whole, and thus environmental suitability worldwide may be overestimated. However, fall armyworm’s wide observed distribution in Africa suggests any overestimation is slight. The predominance of the maize strain in Africa may influence the establishment of fall armyworm from Africa in environmentally suitable parts of Asia, where rice is much more widely grown than maize.

Diet can affect temperature tolerances, and the temperature threshold for development was several degrees lower when fall armyworms were fed leaves from early vegetative maize plants then when fed leaves from late vegetative or reproductive plants^38^ (Supplementary Table S1). Environmental conditions can alter the impacts of biopesticides, and infection rates of diseases and natural enemies that control pests, including in fall armyworm^3,39,40^. Predictions of range, abundance, impacts, and the outcomes of Integrated Pest Management strategies would therefore benefit from a better understanding of the relationship between strains, diet, pesticide effectiveness and environmental limits on distributions. Given the encouragingly robust results of SDMs based on climate and land use variables, future work could extend statistical modelling to the relationship between environmental suitability and fall armyworm abundance and impact on crops. If data on the distribution of potential biocontrol agents could be obtained, their environmental suitability for these species could be also studied using SDMs.

In conclusion, there is considerable potential for near global invasion and seasonal migration of fall armyworm. Vigilance is needed to monitor for the onward invasion of fall armyworm via potential migration routes into North Africa and South Asia, and on some high-risk trade and air travel routes. Management decisions would be improved by further research on fall armyworm’s seasonal migration and population dynamics, and the environmental dependency of interactions with other species.

## Materials and methods

### Effect of abiotic and host plant characteristics on life-cycle

To forecast a species’ potential range, it is necessary to consider the environmental factors that are needed for the species to complete its life cycle. These factors could directly limit the target species’ distribution, and are often termed ‘proximal’ variables. Using these variables increase accuracy and biological realism of projections of species distributions following invasion (also termed ‘transferability’^41^). We reviewed the literature of field observations and experimental studies into fall armyworm to investigate the linkages between the environment and life cycle (Fig. 1, Supplementary Table S1). The studies were conducted on different populations and the population’s strain (which could affect physiology, see below) was rarely reported. Nonetheless, informative patterns emerged. We summarise the patterns below and in Fig. 1, but full details and references are in Supplementary Table S1.

The most commonly studied relationship between life-history stage and environment is the effect of air temperature on larval and pupal survival and development rates. The minimum temperature for development was variously reported as 8.7°C, 13.8°C, 9.5-10.9°C, and 10°C. Several studies find evidence that developmental time of egg, larval, pre-pupal, and pupal stages decreases with temperature up until 32-33.5°C (or even 35°C). However survival of these stages is greatest around 25°C (between 20 and 32°C), and 35°C appears to be an upper limit on survival. Fecundity and adult longevity is greatest between 21 and 25°C. Constant temperatures do not represent real conditions in the field, and studying the effects of fluctuations can be informative. When median temperatures are between 15 and 25°C, daily fluctuations above and below these temperatures increase pupal and larval development rates, and decrease adult deformity.

The importance of moisture and precipitation is complex. Precipitation and irrigation have a direct negative effect on larval and pupal survival. Heavy rainfall fills the maize whorl with water, in which larvae float, until it overflows and the larvae are spilled out or drown (a process which is helped by wind gusts). Rainfall and irrigation are thought to trap moths and drown in their pupation tunnels, with the effects being stronger in more friable soils, when rainfall can also cause the tunnels to collapse. In fact, a local control measurement in Zambia involves spreading ashes in the funnel in order to exacerbate the effect of water (pers comm Patrick Kalama). Lack of moisture during pupal stages appears to have little direct effect on survival or development rates. However, indirect effects of moisture are likely more important for fall armyworm population sizes than direct effects. This is because abundance tends to peak during rainy seasons, particularly in drier sites, possibly because of increased host plant growth. On the other hand, infestation rates are highest in maize deprived of irrigation for the longest, likely because plant moisture stress favours insect development^42^.

There is evidence that maize growth stage positively impacts development speeds (i.e. development is fastest for larvae eating mature leaves). This is unexpected, as leaf nutritional value typically declines with growth stage. It is unclear whether this finding is relevant to field populations, as larvae prefer to feed in whorls on developing leaves, presumably for protection^43^.

There are two genetically distinct fall armyworm ‘strains’, which specialise on maize and rice ^23^. Some inter-strain mating can occur; rice strain females prefer to accept maize strain males, resulting in mixed populations, but maize strain females and rice strain males appear to be reproductively incompatible^20^. Both strains appear to be present in Africa^12,13^. The strains differ in the rates of larval development on the host plants, mating behaviour, use of food resources, resistance to insecticides, and variation in susceptibility to plants expressing *Bacillus thuringiensis* (Bt) proteins^13,20^. There is evidence for variation in the environmental tolerances of the two strains. Basal temperature was found to be 10.6°C and 10.9°C for two populations of the maize strain, and 9.5°C and 9.6°C for two populations of the rice strain (though no statistical test was done)^37^, and seasonal abundance varies differently for the two strains in Florida^44^. There is therefore an argument for constructing separate SDMs for each strain, but the distributions and inter-breeding status of the two strains are not sufficiently known to do this, and we therefore treat the strains as a single entity.

### Environmental data

Based on the life-history and environmental requirements of fall armyworm, and in light of the climatic differences between the Americas and the rest of the world, we selected the following climatic variables:

- SumWet, total amount of precipitation in wettest three months of year (intensity of rainy season, when most food available and population growth is fastest)
- LenWet, number of months when rain is greater than average (length of rainy season, when most food is available and population growth is fastest)
- SeasPpn, seasonality of precipitation (difference in rainfall between rainy and dry season)
- MinTemp, mean temperature of the coldest month of the year (the lowest limit for growth)

We also initially used GDD13.8, annual growing degree days above a lower development threshold of 13.8° C (minimum temperature for survival) was also initially used. We selected 13.8°C as the lower development threshold^45^, as this value is widely cited by other researchers. However, the correlation between MinTemp and GDD13.8 is 0.98. This degree of collinearity can cause inaccurate measurement of the relationship between explanatory and response variable^46^. We removed GDD13.8, as MinTemp has the more intuitive link with species distributions, and a clear temperature threshold for fall armyworm survival is widely reported (Supplementary Table S1). Additionally, fall armyworm populations in west Argentina occur in areas with substantially fewer growing degree days than the populations in the USA. This suggests that the cold range margin is more likely to be set by M¡nTemp than GDD. There is considerable variation in the rainy season between America and other parts of the world, particularly Africa^47^. The precipitation variables were therefore chosen to accommodate variation in the timing and length of the rainy season, or even multiple rainy seasons (e.g. West vs. East Africa). Climatic variables were calculated from monthly averages for the period 1961-1990, derived from the climatic research unit (CRU) dataset at 10 arc-minute resolution^48^.

In addition to climatic variables we also used ‘Forest’, the proportion of each 10 arc-minute grid-cell that is covered by trees. This is because fall armyworm is only reported from agricultural areas, though there may be many areas covered by forest that are climatically suitable for the species, but from which it is not reported or is absent due to a lack of host plant. We used forest rather than crop or pasture land as forest is relatively easier to delineate than grassland using satellite data. Forest cover was drawn from the European Space Agency’s Global Land Cover 2000 project at 1km (<u>https://www.esa-landcover-cci.org/</u>).

### Distribution data

Presence records for the Americas were obtained from three sources. 1) Global Biodiversity Information Facility (ww.gbif.org) in November 2016. Records that did not have coordinates but did have location descriptions were georeferenced with accuracy equal to the climatic grid data. 2) Review of literature on Fall Armyworm in the region. 3) Consultation between CABI and local experts in several countries. In the southern USA, occurrences south of 27 degrees in Florida and south of 31 degrees in Texas were considered to be year-round populations and were included as presence data points. Other USA populations were not included as they represent seasonal migratory populations. In total, 876 presence locations were found. Data and sources for the Americas that can be shared publically are available in Supplementary Table S2. Distribution data for Africa were obtained from four sources. 1) A survey of farming households in Ghana and Zambia, conducted in July 2017^4^. The countries were stratified into geographic regions, within which survey locations were chosen randomly. Surveys yielded 466 incidences of farming households observing symptoms or larvae of fall armyworm in their fields. 2) Published literature^13,49^. 3) Infestations of fall armyworm reported to CABI’s Plantwise clinics in Ghana and Zambia from July 2016 – June 2017. 4) Seven pheromone traps managed by the USAID Agricultural Development and Value Chain Enhancement (ADVANCE) project, from April 2017 - July 2017.

Distribution data were filtered so that only one presence was recorded in each climatic grid-cell, resulting in 240 presences in Africa, and 167 presences in the Americas. Due to the recent dedicated searches, the African distribution was better sampled than the American distribution. The difference in sampling intensity between the two continents led to concerns that SDMs might be over-fitted to the well-studied locations in Ghana and Zambia. This would underestimate the suitability of areas outside Ghana and Zambia, particularly areas that are environmentally similar to the native range, but different to the Ghanaian and Zambian range. We therefore sub-sampled several proportions of the African distribution (5, 10, 20, 30, 50, and 70%) and used four sub-samples at each proportion to construct alternative SDMs.

In order to select pseudo-absences, we used two approaches to delimit the geographic background in the Americas and Africa to which fall armyworm could reasonably be expected to disperse without human assistance (Figure 2). First, we used all countries in America and Africa in which fall armyworm records were available. American countries were excluded from the backgrounds if they did not have records of fall armyworm but (i) fall armyworm is known to be present (determined using CABI’s Crop Protection Compendium, <u>https://www.cabi.org/cpc/about/</u>, and internet searches), or (ii) if the country is surrounded by countries in which fall armyworm is recorded. This exclusion avoided the inclusion of countries in the background where the species is present but has not been sampled, reducing the probability of ‘false absences’. Second, we investigated the impacts of restricting the geographic background to the region within 500km of presence points (Fig. 2). Specifying an upper distance can help prevent models from contrasting completely different climate conditions, e.g. temperate vs. tropical^50^. This contrast would only yield the information that fall armyworm can live in the tropics all year round. For both approaches, pseudo-absences were randomly placed in climatic grid-cells within the background region, but outside occupied grid-cells. The placement of pseudo-absences was repeated 20 times for each sub-sample. In each distribution dataset, the number of pseudo-absences was the same as the number of presences used.

In total, we ran models with 488 datasets: six different levels of sub-sampling of African presences from the entire background (5, 10, 20, 30, 50 and 70%) each with 20 repetitions of sub-sampling, one dataset of all presences using the entire background, and one with the geographically restricted background, and for each of which we randomly sampled pseudo-absences four times.

### Species distribution modelling

An ensemble SDM^51^ was created, which included eight modelling techniques: artificial neural networks (ANN), classification tree analysis (CTA), flexible discriminant analysis (FDA), generalized additive models (GAM), generalized linear models (GLM), multivariate adaptive regression splines (MARS), random forest (RF), and surface range envelope (SRE, note this does not use pseudoabsence data in model calibration but does in validation).

We used internal validation to evaluate SDM accuracy, splitting each of the 488 distribution datasets randomly so that 70% of the presence and pseudo-absence points were used to calibrate the models. These models were used to predict suitability at the 30% remaining validation distribution data points. AUC and TSS were used to judge how accurately the models predicted the validation data.

For each of the distribution datasets we constructed an ensemble forecast for the global terrestrial surface. Ensembles were made using models (from the 488 distribution datasets) for which validation TSS≥0.4. Models with TSS≥0.4 are considered to have ‘moderate’ performance (greater than ‘fair’, but less than ‘substantial’^52^). However, we used all presence and pseudo-absence points in the given dataset to construct the final models to be included in the ensemble (i.e. not just the 70% of the data used in calibration). This was to ensure that when a combination of dataset and technique yielded moderate accuracy, all of the data in that dataset were then used to maximise the information in the final model. In order to be confident that TSS and AUCs of the internally validated models reflected the accuracy of the full models used to project distributions, we calculated the Spearman’s correlation between global projections from internally validated and full models. To construct ensembles, the selected models were rescaled so that projections were on the same numerical scale, and the mean suitability predicted by all retained models was calculated, weighted by the accuracy (TSS) of each model. This method has been shown to be the most accurate of the ‘traditional’ ensembling methods that could be applied to these data^53^.

In order to investigate the effect of biased recorder effort and geographic background on environmental suitability, we compared the agreement of the global ensemble projections made using each distribution dataset using Cohen’s Kappa and balanced accuracy^54^. For the latter, we binned projected suitabilities into 20 percentile bins, and calculated how well the projections from one dataset classified the projections from another dataset.

In order to determine whether the environment in the geographic region from which distribution data were drawn is representative of the entire global terrestrial surface we calculated the Multivariate Environmental Similarity Surface^55, 56^. If there are environmental conditions somewhere in the world that have “no analogue” with environmental conditions in the American and African backgrounds, SDM projections into these regions would be extrapolations, with added uncertainty.

The importance of environmental variables for fall armyworm’s range was calculated using all of the distribution data in a given dataset, and using all models, regardless of TSS score. For any given environmental variable, that variable was randomised, an SDM was made with the shuffled dataset, and the Pearson’s correlation (r) calculated between the SDMs with original and shuffled data. Importance is calculated as 1-r, so a value 0 indicates the variable has no influence on the SDM.

### Has the fall armyworm invaded where we expect it to, based on the native distribution in the Americas?

To answer this we calculated the niche expansion between the Americas and Africa, using the methodology developed by Broennimann, et al.^57^. The environment (climate and forest cover) in the two regions is decomposed into the most important trends using a Principal Components Analysis (PCA). We used all distribution data from the native year-round range, and from the studied African range in Fig. 2 (i.e. there was no sub-sampling). We then divided this environmental space into a grid of 100 × 100 cells. We measured the density of species occurrences for each combination of environmental conditions in each grid-cell of the environmental space using a kernel smoother function to correct for sampling bias and environmental availability and to ensure that the results were independent of the grid resolution. These analyses were done using ecospat package^58^ in R v3.4.1^59^. We used these gridded data to calculate how much of the African niche remained within the native niche (‘niche stability’). Niche stability is the proportion of the densities in the colonised range that overlaps with the native range.

### Transportation beyond Africa

In order to illustrate the potential for fall armyworm to spread from Africa to other parts of the world, we first identified the countries most likely to act as sources for fall armyworm, and those most vulnerable to fall armyworm establishment, as those with >33660km^2^ of suitable climate (i.e. 100 10 arc-minute grid-cells with climate suitability >0.5). This resulted in 64 countries being identified as sources or vulnerable. We then examined two major pathways for invertebrate introduction: trade and passenger air travel. Trade is one of the main drivers of plant pest introduction globally^10^. We used values of all traded goods, as the propensity of fall armyworm to be transported with particular commodity types is not yet known, and all trade provides a reasonable estimate of introduction likelihood for a range of species^36^. We obtained United Nations data on total exports from sub-Saharan African countries to all countries for the period 2012-2016 (2017 data appeared too incomplete to use). Data were obtained from the UN Comtrade database (<u>https://comtrade.un.org/</u>). We considered the trade routes most likely to transport fall armyworm as those with a total trade volume >500,000,000 USD during the reporting period (the top 5% of trade routes from source to vulnerable countries). Passenger air travel is suggested to be the route by which fall armyworm was first introduced to Africa, and is thought to be important in insect introductions^29^. We used data on the number of passengers in 2013 whose embarkation point is in a sub-Saharan African country, and whose final destination is outside sub-Saharan Africa. Data were obtained from the VBD-A¡r tool^60^. We considered the country to country air travel routes most likely to transport fall armyworm as those that carry >10000 passengers during the reporting period (the top 13% of air travel routes from source to vulnerable countries). We then mapped the trade and air travel routes from source countries in sub-Saharan Africa to vulnerable countries worldwide.

### Data availability statement

Some distribution data from South America analysed during this study are included in the Supplementary Information files. This does not include data from Plantwise clinics in Bolivia, Honduras, Nicaragua and Peru, due to data sharing restrictions. Some other distribution data are available from CABI’s Plantwise programme but restrictions apply to the availability of these data, which were used under license for the current study, and so are not publicly available. Data may be available from the authors upon reasonable request and with permission of Plantwise. All other data used are publically available from the referenced data sources.

## Acknowledgements

We are very grateful to Stephanie Wheeler for georeferencing, Dean Paini for information on fall armyworm introduction likelihood, and Jason Chapman for discussion on migration. The authors gratefully acknowledge the financial support of the UK Department for International Development (DFID) and PRISE project (UKSA IPP Call 1). We wish to acknowledge the support of our Plantwise donors: DFID (UK), SDC (Switzerland), DEVCO (European Commission), DGIS (Netherlands), IFAD, Irish Aid and ACIAR (Australia). We would also the like to thank the Ministry of Food & Agriculture, Ghana and Ministry of Agriculture and Livestock and Ministry of Local Government and Rural Development, Zambia, for the extension service providers in their function of plant doctors for gathering plant clinic data and the in-country plant clinic data managers for managing and uploading the data. CABI gratefully acknowledge the core financial support from our member countries (and lead agencies) including the United Kingdom (Department for International Development), China (Chinese Ministry of Agriculture), Australia (Australian Centre for International Agricultural Research), Canada (Agriculture and Agri-Food Canada), Netherlands (Directorate-General for International Cooperation), and Switzerland (Swiss Agency for Development and Cooperation).

## Author Contributions

RE conceived the study, collected data, performed analysis, and drafted the paper. PG and SM contributed to the acquisition and interpretation of data for the project, and revised the manuscript.

## Competing Interests Statement

The authors declare no competing interests.

## List of supplementary material

Table S1. Summary of evidence for fall armyworm developmental and population responses to the environment extracted from literature sources.

Table S2. Distribution data from the Americas

Figure S1. Effect of different sub-sampling proportions and pseudo-absence selection diameters on model predictions (maps)

Figure S2. Effect of different sub-sampling proportions and pseudo-absence selection diameters on Balanced Accuracy

Figure S3. Histograms of each environmental variable in 10 arc-minute grid-cells from which the fall armyworm is recorded.

Figure S4. Multivariate Environmental Similarity Surface analysis.

